# Unbreakable DNA tension probes show that cell adhesion receptors detect the molecular force-extension curve of their ligands

**DOI:** 10.1101/2022.04.04.487040

**Authors:** Rachel L. Bender, Hiroaki Ogasawara, Anna V. Kellner, Arventh Velusamy, Khalid Salaita

## Abstract

Integrin receptors transduce the mechanical properties of the extracellular matrix. Past studies using DNA probes showed that integrins sense the magnitude of ligand forces with pN resolution. An open question is whether integrin receptors also sense the force-extension trajectory of their ligands. The challenge in addressing this question pertains to the lack of molecular probes that can control force-extension trajectories independently of force magnitude. To address this limitation, we synthesized two reversible DNA probes that fold with identical self-complementary domains but with different topologies. Thus, these probes unfold at the same steady-state force magnitude but following different kinetic pathways to reach the fully extended ssDNA state. Hairpin-like probes unzip with a low barrier of 14 pN while the pseudo-knot-like probes shear at 59 pN. Confirming that we had created probes with different barriers of unfolding, we quantified platelet integrin forces and measured 50-fold more tension signal with the unzipping probes over the shearing probes. In contrast, fibroblasts opened both probes to similar levels indicating more static forces. Surprisingly, fibroblast mechanotransduction markers, such as YAP levels, fibronectin production, actin organization, and integrin activation were significantly elevated on unzipping probes. This demonstrates that integrin receptors within focal adhesions sense the molecular force-extension profile of their ligands and not only the magnitude of equilibrium mechanical resistance.

## Introduction

Integrin receptors are heterodimeric transmembrane proteins responsible for linking the internal cytoskeleton of the cell with the outside extracellular matrix (ECM).^1, 2^ Because integrins transmit cell-generated forces to the ECM, it is not surprising that this class of receptors are mechanotransducers. For example, binding of talin to the cytoplasmic tail of integrins leads to “inside-out” activation and involves a conformational shift of the integrin to the open extended state that has a ∼ 1000 fold enhancement in affinity toward the ECM.^3, 4, 5, 6^ Conversely, “outside-in” signaling requires that integrins bind clustered and mechanically stable ECM ligands that resist traction forces of tens of piconewtons (pNs) per molecule to trigger activation.^7^ Integrins also tune the applied forces transmitted to the ECM ligands in response to the mechanical properties of the ECM itself. ^8, 9, 10, 11^ This is illustrated by experiments showing that cells cultured on stiff polyacrylamide gels generate greater traction forces than cells cultured on low modulus gels.^12, 13^ Cells also test ECM rigidity with high spatial resolution as evidenced by their transient deflection of 500 nm PDMS micropillars.^14^ Integrin mechanosensing at molecular scales has been studied using a suite of nucleic acid probes that include DNA duplexes that rupture at specific mechanical thresholds.^15^ These probes can be designed in unzipping and shearing conformations with differential barriers to rupture, estimated at ∼ 12 and ∼ 56 pN, respectively, assuming a 2 second force duration. These probes are useful in manipulating the maximum integrin tension and recording the ensuing cell signaling state.^16, 17, 18, 19, 20^ Indeed, previous studies indicated that > 43 pN forces are required for initial cell adhesion.^15, 21^ Because DNA duplex probes rupture, mechanotransduction is terminated and hence such probes are poorly suited for measurement of forces. Recently, the Liu group constructed a DNA-based reversible shearing probe in an attempt to overcome this limitation.^22^ Using this reversible probe and other innovative tension sensor designs a more clear description of the forces associated with integrin activation have emerged indicating that *F* > 56 pN are central to focal adhesion maturation. ^11, 23, 24, 25^

Interestingly, other studies have suggested that mechanosensor proteins not only detect and transduce force magnitude but also the force-extension history upon engaging the ligand.^26, 27^ For example, mechanical strain of integrin-ligand bonds leads to their reinforcement which is described as catch-bond behavior.^28, 29^ Integrin-ligand bonds also respond to pN mechanical cycling by transitioning to a long-lived binding state to their ligands.^30^ Therefore, an emerging fundamental question in the field is whether integrins detect the molecular extension profile of ECM ligands in addition to the equilibrium resistive force applied by the ECM. Addressing this question poses as an experimental challenge as it requires developing molecular probes that offer identical equilibrium responses to tension but that unfold with unique barriers and hence differing mechanical force-extension responses. In other words, the problem is that current probes are designed to measure or manipulate equilibrium force magnitude rather than the force-extension curve. ^24, 31, 32^

To address this challenge, we generated two DNA probes that form folded structures due to self-complementary domains. The reversible unzipping (RU) probe adopts a classic stem-loop hairpin while the reversible shearing (RS) probe was synthesized to fold into a pseudo-knot like structure using a 3’-3’ linkage **(Fig. 1a)**. Because the nucleobase composition is identical, both DNA probes display identical *F*_eq_ values (the equilibrium force that leads to a 50% probability of unfolding), yet with different kinetic barriers to unfolding and different force-extension curves. In this manner, these probes diverge in their energy landscapes toward mechanical unfolding but display identical thermal unfolding response. Probes were covalently anchored to a surface at one terminus and displayed integrin ligands at the second terminus. Thus, when cells are seeded to the surface, integrins receptors will bind to the probes and transmit forces that lead to unfolding at unique magnitudes of tension **(Fig. 1b)**. The rationale for the work was that RU and RS probes would provide insights into the dynamics of integrin-transmitted forces as RS and RU probes should show similar signal if forces are stable (equilibrium), in contrast to dynamic forces or transient forces that would preferentially unfold the RU probes over the RS probes.

**Figure 1.**
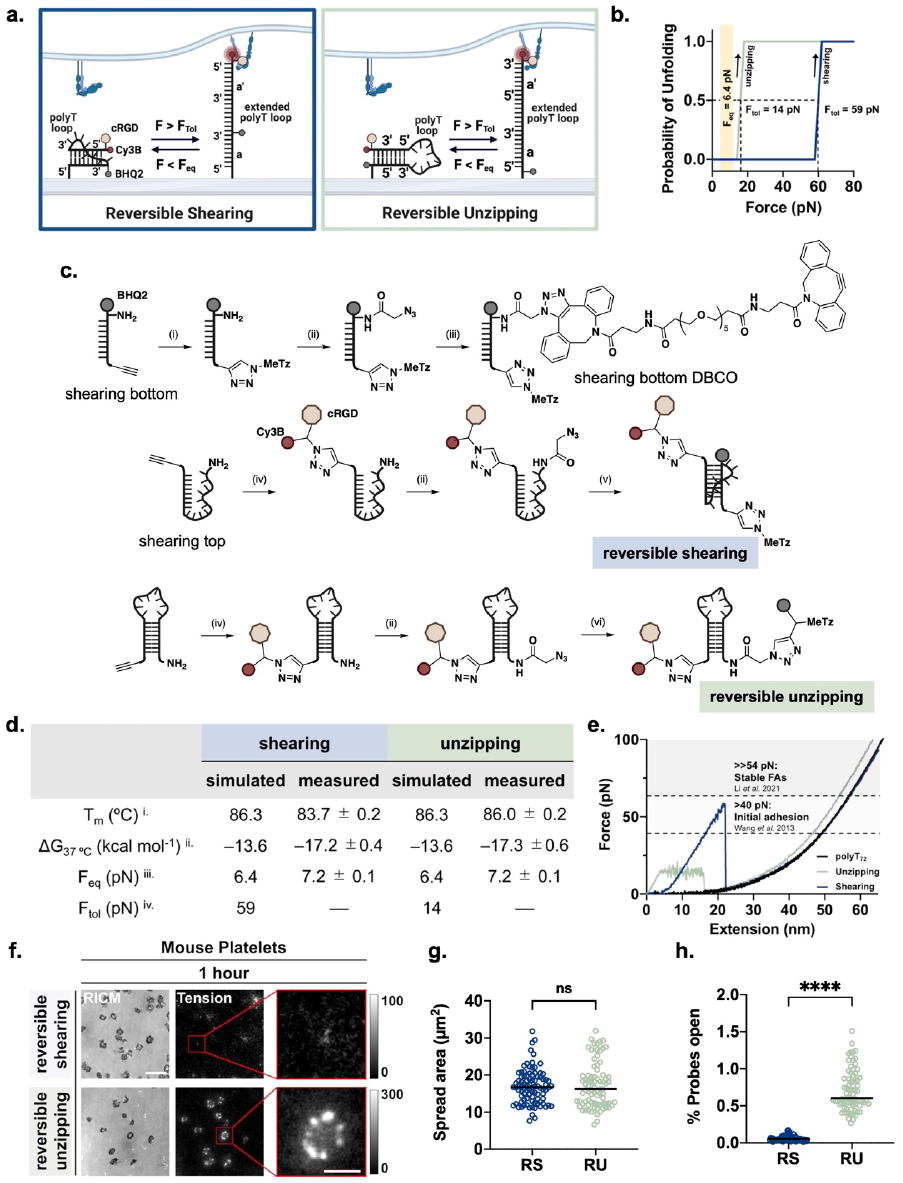
Design and characterization of RS and RU probes. **a**. Scheme depicting RS and RU probes immobilized on a glass surface. **b**. Plot of probability of RS (blue) and RU (green) probe unfolding at increasing force. Data generated from oxDNA simulation (see **SI note 2**). The yellow vertical box corresponds to the *F*_eq_ which was inferred from NUPAK calculations. The vertical dashed lines correspond to the *F*_tol_ of the RS and RU probes at 59 pN and 14 pN, respectively. **c**. Scheme depicting synthesis of RS and RU probes. i) methyltetrazine-PEG_4_-azide, CuSO_4_, THPTA, sodium ascorbate in 40% of DMSO and 60% of water at 50ºC for 1 hour. ii) Azido-NHS ester in aqueous solution of 0.1M sodium bicarbonate and 1X PBS with 20% DMSO at room temperature (r.t.) for 1 hour. iii) DBCO-PEG_5_-DBCO in 50% of water and 50% of DMSO at r.t. for 1 hour. iv) cRGD-Cy3B-N_3_, CuSO_4_, THPTA, sodium ascorbate in 40% of DMSO and 60% of water at 50ºC for 1 hour. v) MeTz-BHQ2-alkyne, CuSO_4_, THPTA, sodium ascorbate in 60% of DMSO and 40% of water at 50ºC for 1 hour. vi) RS-bottom-DBCO in water at r.t. for 4 hours. **d**. Thermodynamic properties, *F*_eq_, and *F*_tol_ of RS and RU probes. i. Tm were measured experimentally in 1X PBS at 10 nM ii. *Δ*G values were calculated from a van’t Hoff analysis obtained from fluorescent melting curves *(see* ***SI note 1***) iii. *F*_eq_ values were calculated based on *Δ*G_folding_.^36^ iv. *F*_tol_ values were simulated using oxDNA software *(see* ***SI note 2***). **e**. Force-extension curve of RS (blue) and RU (green) probes superimposed on the force extension curve of a 72 nucleotide polyT DNA strand (black). Arrows on the graph indicate force induced dissociation of the duplexes, either by shearing or unzipping. The graph also indicates the force threshold required for initial cell adhesion, light grey (> 43 pN), and focal adhesion formation force in dark grey (>> 54 pN). **f**. RICM and fluorescent tension images of mouse platelets seeded on RS and RU probes for ∼ 1 hour. Scale bar, 2 µm. **g**. Plots showing spread area of mouse platelets on RS and RU probes. Spread area was measured by drawing a region of interest around platelets in the RICM channel (*n* = 3 experiments, RS = 90 cells, RU = 81 cells; *p* = 0.5889). **h**. Percent of RS and RU probes opened by mouse platelets. Percent of probes open was determined by dividing the fluorescent tension signal by the fluorescence value of an unquenched surface (*n* = 3 experiments, RS = 72 cells, RU = 77 cells; *p* < 0.0001).

First we performed van’t Hoff analysis and confirmed that RS and RU probes show identical thermodynamic parameters for thermal melting, which indicates that the *F*_eq_ is similar for both probes at 7.2 pN, in good agreement with the NUPAK estimation of 6.4 pN. In contrast, the force tolerance (*F*_tol_), which we define as the peak force required to overcome the barrier to unfolding, was 59 pN and 14 pN for RS and RU probes, respectively, as determined by coarse grain (xDNA) modeling. We found that mouse platelet integrins mechanically unfold the RU probes but do not sustain a sufficient magnitude and duration of force to open the RS probes. In contrast, fibroblast integrins unfold both the RS and RU probes to similar levels after 1 hour of being seeded on the probes. Based on prior measurements of fibroblast and platelet integrin force magnitudes, these results reveal that the force lifetime of the platelet integrin α_IIb_β_3_-cyclic RGD interaction is smaller than the fibroblast integrins α_5_β_1_- and α_V_β_3_-cyclic RGD force duration. Counterinuitively, we found that after 3 hours of cell culture, fibroblasts cultured on RU probes opened fewer probes than when cultured on RS probes. Moreover, fibroblasts on RU probes displayed enhanced mechanotransduction markers such as actin stress fiber formation and nuclear YAP localization, as well elevated levels of activated integrins and ECM secretion. Measurement of integrin tension turnover revealed that integrin-ECM complexes were more dynamic in fibroblasts cultured on RS probes compared to the RU probes which is a consequence of differential mechanical signaling. Because RS and RU probes unfolding is identical at equilibrium force, these results show that integrin adhesion receptors distinguish and transduce the molecular force-extension curves of their ligands and the abrupt molecular extension of RS probes likely hinders mechanotransduction. This work therefore reveals the dynamic aspects of chemo-mechanical signaling in cells.

## Results

### Synthesis and Design of Reversible Probes

The RS and RU probes are comprised of self-complementary DNA sequences linked by a 30 nt polyT spacer and covalently anchored to a surface. The RU probe is a conventional 3’–5’ polynucleic acid that forms a hairpin, while the RS probe is designed with an identical sequence but incorporating a 3’–3’ linkage. We covalently functionalized the RS and RU probes with a Cy3B fluorophore and BHQ2 quencher, enabling the use of fluorescence as a readout for cell-mediated DNA unfolding and traction force. Given that the probe design is completely covalent in nature, only mechanical separation exceeding the *F*_tol_ of the probe leads to an increase in fluorescence.

The RS and RU probes required the introduction of two chromophores, a cyclic peptide, and a tetrazine moiety to drive an inverse electron demand Diels-Alder reaction for surface immobilization. Moreover, the 3’-3’ linkage of the RS probe is not accessible by enzymatic or solid-phase synthesis. Hence, a multistep strategy was required to generate the constructs which were synthesized as shown in **Fig. 1c** and **Scheme S1**. The RU probe and the top strand of the RS probe were first covalently conjugated to the fluorophore labelled ligand using a copper-mediated click reaction (CUAAC), then functionalized with an azide using an NHS ester reaction. Similarly, CUAAC was used to functionalize the quencher-labelled bottom strand with a tetrazine moiety used for surface attachment to the trans-cyclooctene surface, and then functionalized with an azide using an NHS ester reaction. The azide on the unzipping probe was used to functionalize the DNA with the quencher-tetrazine moiety via CUAAC. The azide-functionalized 3’ ends of the top and bottom strands of the RS probe were covalently attached to each other using subsequent strain-promoted click reactions with a homobifunctional DBCO linker. The final products were purified using HPLC and validated by ESI-MS (**Fig. S2** and **S3**).

### RU and RS probes have identical ΔG’s of unfolding but differ in their peak unfolding force

We next measured the *Δ*G of folding for the RU and RS probes. In this study, the temperature-dependent fluorescence of Cy3B was measured from 37 to 95 ºC, allowing us to detect thermally-driven probe unfolding. The melting curves were used to generate a van’t Hoff plot (**Fig. S4, Table S2, S3, SI Note 1**).^33^ As we expected, the thermodynamic parameters of the RS and RU probes were similar. Specifically, the *Δ*G_folding_ (T = 37 ºC) for the RS and RU probes were -17.2 ± 0.4 and -17.3 ± 0.6 kcal mol^-1^, respectively (**Fig. 1d**). These values were then used to infer the *F*_eq_ for the RS and RU probes.^34, 35^

Next, using a detailed coarse-grained DNA model (oxDNA) we simulated force extension curves for the reversible probes (**Fig. 1e**). The dissociation behavior observed for the probes was similar to the behavior observed for corresponding DNA duplex unzipping and shearing (**Fig. S5**). While the RU probe required low levels of force to unfold (*F*_tol_ ∼ 14 pN), the RS probe required ∼ 59 pN of force to dissociate. In both the RU and RS probes, we observed a “two-state” mechanism of unfolding in which the probe’s *F*_tol_ is required for initial unzipping or shearing and is followed by an approximately 25 nm extension of the probe which occurs at a significantly lower force than the probe’s *F*_tol_. As the probes reach full extension, the amount of force the probes can withstand increases into the nN range due to their covalent linkage which prevents rupture.

### Platelet integrin forces unfold RU but not RS probes

As an initial model, we first cultured platelets on RS and RU probes. Platelets are a useful model because of their small size and their ability to generate force.^34, 36, 37, 38, 39, 40^ Understanding platelet force is also biomedically relevant in the study of clotting disorders, wound healing, and drug interactions for common cardiovascular diseases.^41, 42^ Platelets were cultured on RS and RU probe for 1 hour before activation with 10 µM ADP for 10 minutes (**Fig. 1f**). Upon activation, platelets rapidly adhered to the surface and generated molecular traction forces. The spread area of platelets cultured on RU and RS probes was similar; however, there was a stark contrast in the fluorescent tension signal (**Fig. 1g**). Although the probes are chemically and thermodynamically identical, platelets cultured on RU probes generated nearly 50-times the amount of fluorescent tension signal compared to platelets cultured on RS probes (**Fig. 1h**). Consistent with the data obtained from oxDNA modeling, this result demonstrates that different force thresholds are required to unfold RS and RU probes. If platelets integrins applied steady state forces that allowed the RS probes to reach their equilibrium unfolded state, then both probes would unfold to similar levels. Hence, the lack of tension signal on the RS probe shows that platelet integrin forces transient when compared to the longer time scales of for nucleic acid mechanical shearing of > 2-30 seconds. ^22, 43^

It is important to note that the differences we observed in the platelets cultured on RS and RU probes were not caused by either the density or the accessibility of the ligand on the surface. We validated that the RS and RU probe surface density was similar by confirming that the fluorescence of unquenched surfaces was the same, indicating that the similar amount of ligand is immobilized (**Fig S6**). Furthermore, the radius of gyration of the ligand on the RS probes is ∼ 7 nm, while the radius of ligand gyration on the RU probes is ∼ 2 nm, given the different location of ligand with respect to the location of surface attachment. Therefore, the ligand on the RS probe is not less accessible than the ligand on the RU probe. Hence, the observed differences in fluorescence signal are due to dynamic nature of platelet integrin-RGD forces rather than due to difference in ligand density or ligand accessibility.

### Fibroblast integrins unfold RS and RU probes in a time-dependent fashion

We next studied the behavior of mouse embryonic fibroblasts (MEFs) stably expressing GFP vinculin (MEF-GFP-vinculin) on surfaces coated with RS and RU probes. Fibroblasts are one of the best studied models for integrin mechanotransduction and their ability to generate large traction forces to mechanosense the ECM is well documented.^44^ MEFs were imaged after 1 and 3 hours of seeding to evaluate cell spread area, tension signal, and GFP-vinculin signal **(Fig. 2a)**. At 1 hour, there was not a significant difference in tension signal produced by cells on RS and RU probes (**Fig. 2b**). Unlike platelets, MEFs open both the RS and RU probes, which indicates that integrin-RGD forces transmitted within focal adhesions are sufficient to mediate unfolding of the shearing probes. We next confirmed that probes are reversible and rapidly refold (∼ 1-2 min) upon termination of the cytoskeleton generated forces by treating cells with Latrunculin B, a disruptor of actin polymerization **(Fig. S7)**.^45^ Surprisingly, after 3 hours, MEF cells cultured on the RS probes appeared to maintain a similar level of tension signal while cells on RU probes generated a significantly lower tension signal at 3 hours compared to that of 1 hour. This is unexpected given that the cell spread area on RU and RS probes was similar (**Fig. 2c**). Furthermore, these probes report on integrin mediated tension in real-time (**Fig. 2 d, e, Video S1**), and one would expect that a difference in tension signal between cells on each of the surfaces would be accompanied by a difference in their focal adhesion size. However, we did not observe differences in the size or number of focal adhesions for cells cultured on RS and RU probes (**Fig. 2f-h**). Perhaps the most striking observation was the presence of strong focal adhesions on RU probes after 3 hours of culture despite the significant decrease in tension signal. These unanticipated differences in cell behavior on RS and RU probes led us to further explore the mechanically-induced biochemical response of MEF cells to these probes.

**Figure 2.**
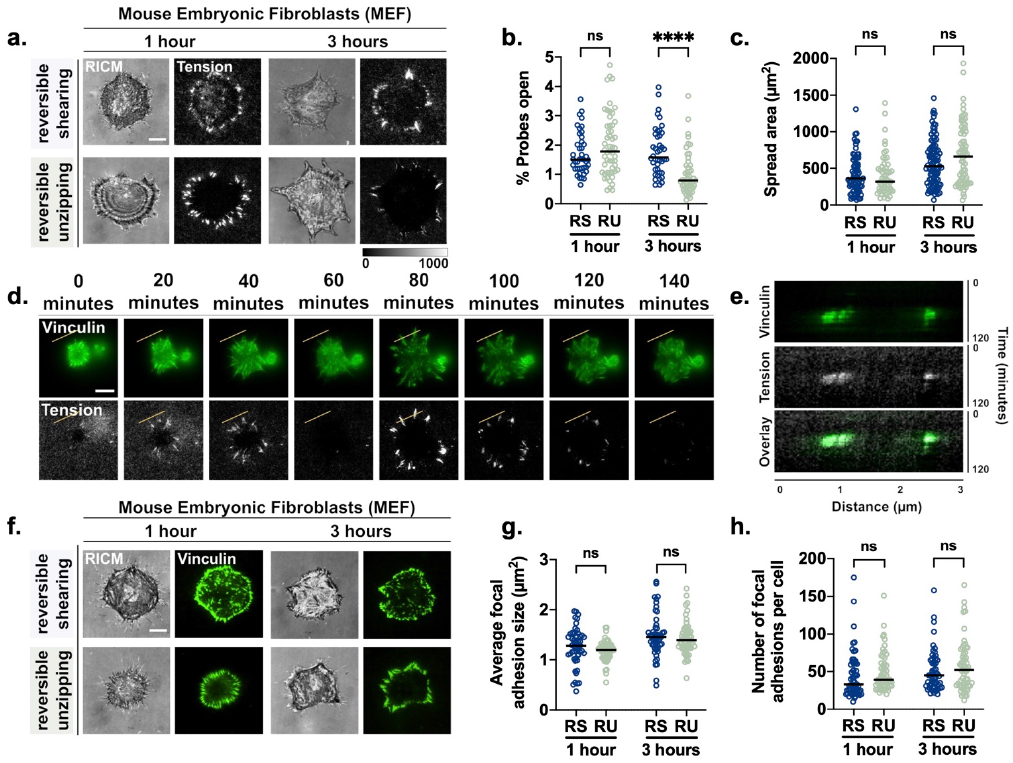
Mouse embryonic fibroblasts (MEFs) respond to RS and RU probes differently. **a**. RICM and fluorescent tension images of MEFs plated on RS and RU probes. **b**. Plots showing %probes open for RS and RU surfaces. Probes open % was determined by normalizing the tension signal against that of an unquenched surface (*n* = 3 experiments, 1 hour: RS = 52 cells, RU = 53 cells, *p* = 0.2106, 3 hours: RS = 52 cells, RU = 50 cells, *p* < 0.0001). **c**. Spread area of MEFs on RS and RU probes (*n* = 3 experiments, 1 hour: RS = 80 cells, RU = 68 cells, *p* = 0.2106, 3 hours: RS = 100 cells, RU = 85 cells, *p* < 0.0001). **d**. Timelapse imaging of vinculin-GFP transfected MEFs on RS probes. **e**. Kymograph from yellow line region of interest in timelapse (d) indicating that vinculin and tension signal colocalize. **f**. RICM and vinculin-GFP images of MEF cells on RS and RU probes. **g**. Quantification of average focal adhesion size in MEFs plated on RS and RU probes (*n* = 3 experiments, 1 hour: RS = 55 cells, RU = 60 cells *p* = 0.4598, 3 hours: RS = 46 cells, RU = 59 cells, *p* = 0.5972). **h**. Quantification of focal adhesion number in MEFs plated on RS and RU probes. Focal adhesion size and number was determined using a particle quantification method in Fiji (**Fig. S8**) (*n* = 3 experiments, 1 hour: RS = 59 cells, RU = 58 cells, *p* = 0.6980, 3 hours: RS = 55 cells, RU = 58 cells, *p* = 0.4051). Tension images were acquired using epifluorescence. Vinculin images were acquired using TIRF, Scale bar, 5 µm.

### Fibroblasts show enhanced mechanical signal on RU probes compared to RS probes

We first chose to investigate actin stress fiber formation and Yes-associated protein (YAP) nuclear translocation as validated markers of cellular mechanotransduction.^46, 47^ Circular actin is associated with early cell adhesion while the progression through radial to linear actin patterns develops as a cell forms stable adhesions to its substrate (**Fig. 3a**). Cells were stained with SiR-actin and subsequent image analysis revealed that cells cultured on RU probes formed greater levels of actin stress fibers compared to cells cultured on RS probes (**Fig. 3b,c**). Actin patterns were further classified as either circular, radial, or linear. Interestingly, actin in cells cultured on RS probes tended to be in the radial form while actin in cells cultured on RU probes was in the linear form (**Fig. 3d**). As a control, we cultured cells on conventional > 56 pN DNA rupture probes that terminate the mechanical signaling pathways. Indeed, cells on these surfaces show minimal stress fiber formation and actin remained primarily in the circular form (**Fig. S9**). This finding suggests that cells cultured on RU probes form more stable adhesions, inconsistent with the tension measurements.

**Figure 3.**
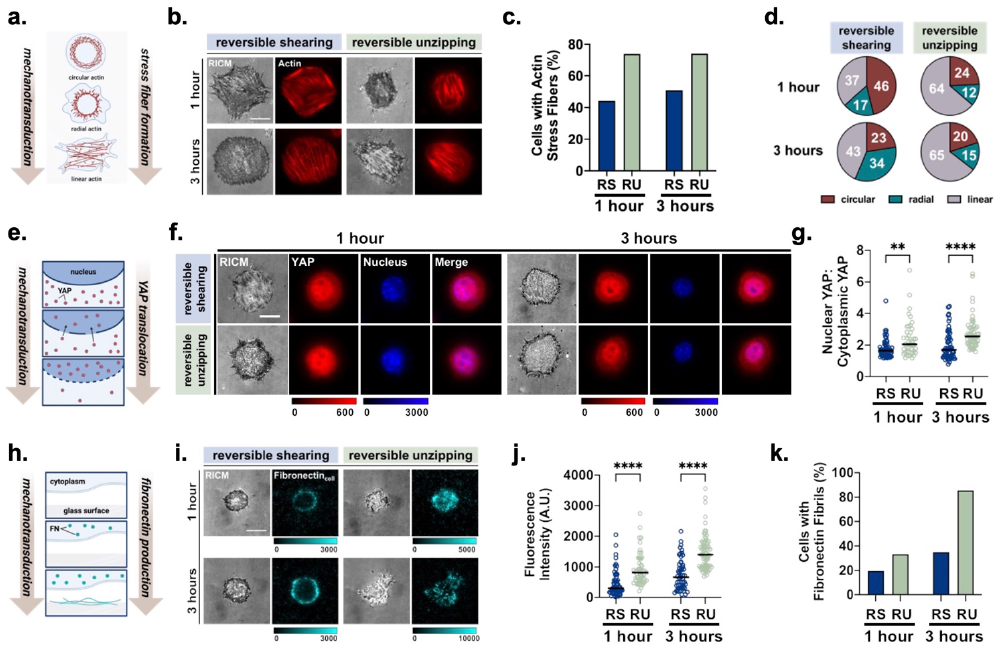
Mechanotransduction markers are greater in MEFs plated on RU probes. **a**. Schematic showing that as cell mechanotransduction increases, stress fibers transition from circular to linear morphology. **b**. RICM and F-actin images (SiR-actin 647) of MEF cells cultured on RS and RU probes for 1 or 3 hours. **c**. Quantification of the percent of MEFs cultured on RS and RU probes that had formed stress fibers (*n* = 3 experiments, 1 hour: RS = 53 cells, RU = 46 cells, 3 hours: RS = 59 cells, RU = 58 cells). **d**. Distribution of actin patterns in MEFs on RS and RU probes (*n* = 3 experiments, 1 hour: RS = 52 cells, RU = 58 cells, 3 hours: RS = 71 cells, RU = 65 cells). **e**. Scheme showing that as cell mechanotransduction increases, nuclear levels of YAP increase. **f**. RICM, YAP, and nuclear staining (DAPI) images for MEF cultured on RU and RS probes for 1 and 3 hours. **g**. Ratio of nuclear to cytoplasmic YAP in MEFs plated on RS and RU probes at 1 and 3 hours (*n* = 3 experiments, 1 hour: RS = 59 cells, RU = 42 cells, *p* = 0.0036, 3 hours: RS = 58 cells, RU = 55 cells, *p* < 0.0001). **h**. Scheme showing that as mechanotransduction increases the amount of fibronectin produced by cells increases. **i**. RICM and fibronectin staining images for MEFs seeded on RS and RU surfaces for 1 and 3 hours. Fibronectin was immunostained and measured using epifluorescence. **j**. Plot quantifying fibronectin generated by cells and deposited onto the surface at 1 and 3 hours (*n* = 3 experiments, RS = 61 cells, RU = 51 cells, *p* < 0.0001, 3 hours: RS = 53 cells, RU = 69 cells, *p* < 0.0001). **k**. Percent of cells plated on RS and RU probes that produced identifiable fibronectin fibrils after 1 and 3 hours (*n* = 3 experiments, 1 hour: RS = 61 cells, RU = 51 cells, 3 hours: RS = 53 cells, RU = 69 cells). Fluorescent regions were determined in i using the ROI generated in the RICM channel. Scale bars, 5 µm.

We next measured the nuclear localization of YAP, a transcription co-activator that is localized to the nucleus of cells in response to mechanical signaling (**Fig. 3e**). Cells were allowed to spread for 1 or 3 hours on RS and RU surfaces coated before fixing and immunostaining for YAP (**Fig. 3f**). We found that at both 1 and 3 hour time points, there was a significantly higher amount of YAP localized to the nucleus of cells cultured on RU probes (**Fig. 3g**). Controls where we cultured cells on irreversible shearing probes showed that YAP remained dispersed in the cytoplasm due to the termination of mechanical signaling in agreement with stress fiber measurements (**Fig. S9**).

We were surprised to find that cells cultured on RU probes had higher markers of mechanotransduction than cells cultured on RS probes, particularly given the observed decrease in their tension signal which would suggest a reduced level of mechanical activity. One possible explanation was that cells cultured on RU probes generate higher levels of their own ECM than cells on RS probes. Fibronectin is one of the primary components of the ECM and is secreted in by cells in response to mechanical signaling (**Fig. 3h**).^48^ Therefore, to test this hypothesis, we immunostained RS and RU surfaces for fibronectin secreted by MEF cells (**Fig. 3i**). Indeed, we found that cells cultured on RU probes produced a greater amount of fibronectin than cells on RS probes at 1 hour after seeding (**Fig. 3j**). Notably, after 3 hours, cells on RU probes had begun secreting fibronectin in distinct fibril-like patterns, unlike cells cultured on the RS probes (**Fig. 3k**). Hence, the increased mechanical signaling in cells cultured on RU probes results in the generation of more fibronectin onto the surface. This surface fibronectin can contribute to the ECM-mediated cell mechanotransduction, interfering with the tension probe-mediated cell mechanotransduction by competing with cRGD for integrin adhesion. Based on this finding, we continued to investigated the differential mechanotransduction markers on RU and RS probes at only the 1 hour time point.

### Fibroblast integrins are more active in cells on RU probes than RS probes

Active integrins depost fibronectin from solution onto their surface. Hence, we next measured if there was a difference in fibronectin retrieved from solution and deposited onto the surface by cells on RU and RS probes by adding fluorescently labeled fibronectin into solution (**Fig. 4a**). Cells displaying a greater number of activated integrins are expected to deposit greater amounts of fibronectin. Interestingly, we found that cells on RU probes retrieved and deposited a significantly greater amount of fibronectin than cells on RS probes (**Fig. 4b**). For cells cultured on RU probes, the tension signal and fluorescent signal from deposited fibronectin are overlapping. This confirms our hypothesis that the fibronectin deposited onto the surface is in direct competition with the cRGD tension probe, providing an explanation for the reduction in tension signal observed in the initial experiment (**Fig. 2a,b**).

**Figure 4.**
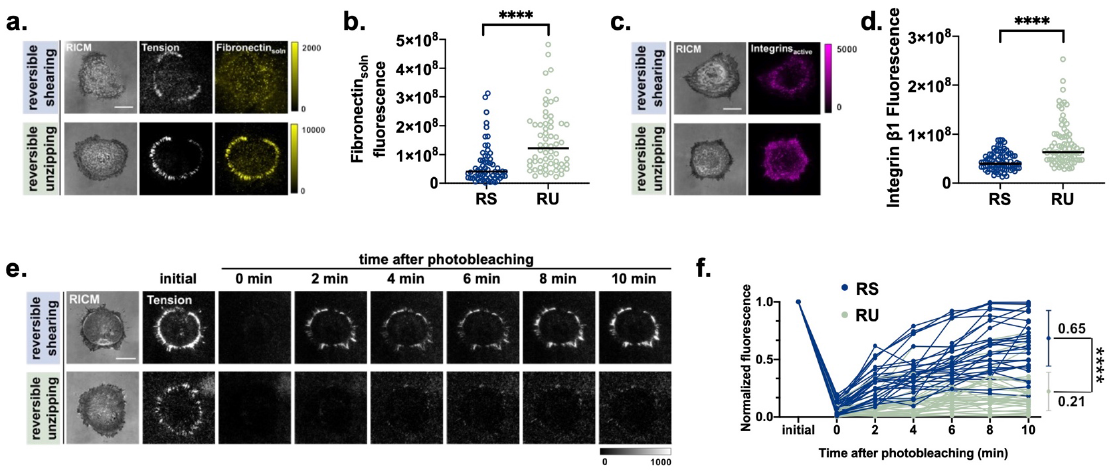
Integrin activation enhanced on RU probes surfaces. **a**. Images showing tension signal and fibronectin retrieved from solution by MEFs plated on RS and RU probes for 1 hour. Fluorescent fibronectin (Alexa488) was added into solution post-plating. Tension was imaged using epifluorescence while fibronectin_soln_ was imaged using TIRF. **b**. Plot of fluorescence of fibronectin retrieved from solution and deposited onto the surface (*n* = 3 experiments, RS = 63 cells, RU = 65 cells *p* < 0.0001). **c**. Epifluorescence immunostaining of active β1 integrins using 9EG7 plated on RS and RU probes for 1 hour. **d**. Plot of active β1 integrin staining per cell on RS and RU probes (*n* = 3 experiments, RS = 86 cells, RU = 80 cells, *p* < 0.0001). **e**. Fluorescent tension signal recovery after photobleaching. Following photobleaching of tension signal, cells plated on RS probes generate new fluorescent tension signal more rapidly than cells plated on RU probes. **f**. Time dependent fluorescent tension signal recovery. Recovery of signal was determined by measuring the fluorescent signal within a region of interest over time. Fibroblasts plated on RS probes recovered 65% of their initial fluorescence while cells plated on RU probes recovered 21% of their fluorescence, demonstrating that the integrin turnover is greater on RS probes (*n* = 3 experiments, RS: 3 regions of interest per cell, 30 regions of interest total, RU: 3 regions of interest per cell, 30 regions of interest total *p* < 0.0001). Fluorescent regions were determined in a and c using the ROI generated in the RICM channel. Scale bars, 5 µm.

We next quantified active integrins on cells cultured on RS vs RU probes using immunostaining methods. Specifically, we detected integrin β1 in the active conformation using the 9EG7 antibody.^49^ Indeed, we found that cells cultured on RU probes had a significantly higher number of integrins in the active conformation, confirming our earlier results and suggesting that integrin activation and the resulting mechanotransduction is driven by the cell’s interaction with their ligand (**Fig. 4 c,d**). As a control, we confirmed that the increase in integrin activation levels was not simply due to an increase in the total number of integrins in cells on RU probes vs RS probes (**Fig. S11**). These results confirmed that the relative total number of integrins on cells on RU and RS probes is the same; further validating the conclusion that integrin activation levels for MEFs is controlled by the probe unfolding trajectory.

Previous literature demonstrates that as integrin activation levels increase, integrin turnover rates decrease.^50^ To test if the observed increase in integrin activation levels in MEFs on RU probes was accompanied by a decrease in their turnover rates, we photobleached the probes under cells and then measured the relative time for fluorescence recovery. In this experiments we selectively photobleached unfolded probes, as the fluorophores in the folded probes are quenched and photo-protected from bleaching. Thus, the recovery of tension signal indicates integrin turnover - either through arrival of new integrins or rebinding of existing integrins to fresh probes sites. We found that cells cultured on RS probes begin to recover their fluorescent signal in < 2 minutes while cells cultured on RU probes take ≥ 10 minutes to begin to recover their fluorescent signal, indicating that cells on cultured RU probes had lower integrin turnover rates and were slower to form new interactions with the ligand (**Fig. 4e, f**). This last result further corroborates the conclusion that integrin mechanotransduction is enhanced on RU probes compared to that of RS probes.

## Discussion

We have synthesized two completely covalent DNA-based probes capable of reporting on cell traction forces in real-time and measuring cell response to unique force extension curves of their ligands. Previously reported DNA-based probes have two primary limitations. One is that they rely on non-covalent interactions, such as biotin-streptavidin, and labile bonds such as thiol-maleimide and thiol-Au interactions, to anchor the probe to the surface which can lead to spontaneous probe dissociation.^51, 52^ Both thiol-maleimide and thiol-Au bonds are particularly labile in cell media which contains mM concentrations of thiols and reducing agents that drive exchange reactions. Our probes are covalently anchored to the surface using a non-exchangeable TCO-Tz bond and are synthesized using covalent chemical conjugation. Second, DNA hairpin probes cannot measure in real-time cell mechanical events with peak forces exceeding 19 pN. While the reversible shearing probes reported by Liu et al. addresses this limitation, these probes use hybridization to introduce the quencher, and hence under sufficient force the quencher labelled strand will undergo irreversible “peeling” or denaturation (**Fig. S5**).^53, 54^ We have synthesized these probes in the unzipping and shearing conformation, with the shearing conformation containing a novel 3’-3’ linkage that is not accessible using enzymatic or conventional solid-phase phosphoramidite chemistry. Our probes are thermodynamically identical but have unique force thresholds and pathways of unfolding which we have demonstrated with modeling and experimental data.

The RS probe reversibly unfolds at ∼ 59 pN while the RU probe reversibly unfolds at ∼ 14 pN. Following unfolding of each duplex, the probes follow the wormlike chain (WLC) model as they are extended. Thus, given the force thresholds of the probes and their previously identified relationship to cell adhesion, the cells cultured on the shearing probe form initial adhesions before dissociating the duplex while cells cultured on the unzipping probe must dissociate their duplex prior to forming initial adhesions. Full extension of both the RS and RU probe structures results in a mechanically robust ligand with a force rupture threshold in the nN range and thus facilitating maturation of focal adhesions to similar levels. It is important to note that the RS and RU probe each follow a unique force-extension curve as they approach full extension. Therefore, we investigated if the pathways of ligand-force-extension have an effect on the pathways of mechanotransduction in fibroblasts and, if in the absence of thermodynamic differences, the geometry and subsequent kinetic behavior of a probe controls cell response.

As previously mentioned, the RS and RU probes are thermodynamically identical. Thus, if platelets applied continuous force, one would expect that both the RS and RU probes would unfold under constant force as described by the Bell model.^55^ However, platelets cultured on RU probes generate a fluorescent signal that is 50-times higher than the signal generated by platelets cultured on RS probes, indicating that platelet generated force is transient **(Fig. 1f,h)**. Given the published rate constants for force-induced shearing of DNA duplexes, it is likely that integrin forces need to be maintained for ∼ 100 seconds at ∼ 20 pN and ∼ 1 second for ∼ 35 pN in order to unfold the RS probe.^43^ In contrast to platelets, fibroblasts were capable of unfolding both the RS and RU probes resulting in the fluorescent tension signal **(Fig 2a,b)**.

MEF cells were capable of exerting forces greater than the *F*_eq_ of both the RS and RU probes and there was not a significant difference in the average spread area or focal adhesion number and size in cells cultured on either probe. However, between 1 and 3 hours, the average tension signal generated by cells on RU probes decreased by nearly 60%, a surprising result considering there was no measurable decrease in their focal adhesion formation. This was accompanied by increases in other markers of mechanotransduction, specifically, an increase in actin stress fiber formation, nuclear YAP localization, and fibronectin production. Cells cultured on RU probes also had a significantly higher number of β1 integrins in the active conformation, as well as lower integrin turnover compared to cells cultured on RS probes. As we described earlier, the differences we observed in the cells cultured on RS and RU probes were not caused by either the density or the accessibility of the ligand on the surface.

Using RS and RU probes, we have uncovered a unique biological feature of adhesion receptors. Our results suggest that adhesion receptors are not only able to detect the relative force thresholds of their ligands but also their unique force-extension curves. Indeed, prior work has demonstrated that integrins are sensitive to more than just the magnitude of force being applied.^56^ For example, it is known that integrin activation is also sensitive to the force loading history and the direction of force.^57, 58^ Furthermore, integrin activation is known to mediate mechanotransduction within the cell, which in turn influeneces integrin activation on the outside of the cell, highlighting the relationship between extracellular and intracellular signaling.^59, 60, 61^

Previous work has demonstrated that cells require ∼ 43 pN of force to form initial adhesions and greater than 56 pN of force to form stable focal adhesions (**Fig. 5a**). Due to the nature of unfolding of the RS and RU probes, cells cultured on these probes reach these force thresholds by different mechanisms. The *F*_tol_ of the RS probe is ∼ 59 pN while the *F*_tol_ of the RU probe is ∼ 14 pN, assuming a loading rate consistent with that used for modeling. Hence, cells can form initial adhesions on the RS probe before the DNA duplex is sheared. However, after the probe is sheared, the probe unfolds, dropping the force to zero and causing a disruption in mechanotransduction that significantly affects focal adhesion maturation (**Fig. 5b**). Conversely, cells on RU probes cannot form initial adhesions until the probe is fully extended. Therefore, there is no disruption in mechanotransduction as the adhesions mature, possibly explaining the increased markers of mechanotransduction (**Fig. 5c**).

**Figure 5.**
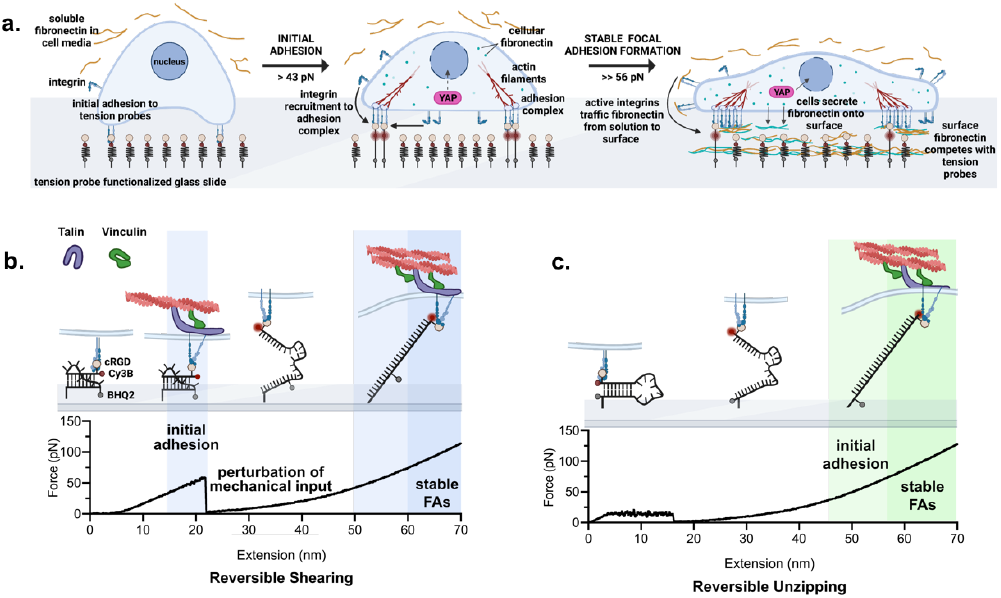
Suggested mechanism for differential response to RU and RS probes by adhesion receptors. **a**. Cells are added to the surface and make initial contact with surface immobilized tension probes. Initial adhesions are formed after the cell exerts > 43 pN of force on the probe. This begins a signaling cascade that leads to an increase in mechanotransduction markers including integrin recruitment to the adhesion complex, YAP translocation to the nucleus, the production of fibronectin by the cell, and the formation of actin stress fibers. Following the cell exerting >> 56 pN of force on the probe, there is further increase in mechanotransduction markers, leading to maturation of focal adhesions, increased YAP translocation to the nucleus, activation of β1 integrins, surface deposition of cellular fibronectin, and retrieval and surface deposition of fibronectin from solution by active integrins. **b**. Force extension behavior of RS probes as it relates to focal adhesion complex formation. The rupture of the DNA duplex in the RS probe requires > 59 pN of force therefore cells on RS probes form initial adhesions as they begin to pull on the probe. Following rupture of the duplex, the resistive force from the probe on the integrin drops to zero, terminating mechanical input. The integrin then fully extends the probe to its contour length, at which point the resistive force is >> 56 pN, and therefore sufficient force for the cell to form stable focal adhesions. **c**. Force extension behavior of RU probes as it relates to focal adhesion complex formation. The rupture of the DNA duplex in the RU probe requires only 14 pN of force therefore cells on RU probes cannot form initial adhesions until the integrin has fully extended the probe to its contour length. Therefore, there is no disruption in mechanical input as the probe is unfolded.

The catch-bond behavior of integrins-ligand bonds may offer a mechanism to explain the unusual response to RS probes. For example, it is plausible that integrins will dissociate from the RS probe after initial shearing, as this transition involves dropping the resistive force from 60 pN to 0 pN, allowing the integrin to rapidly dissociate from its ligand and thus resulting in the perturbation of mechanotransduction. Additionally, the RS probe requires a full extension to its contour length of ∼ 20 nm (based on simulation using oxDNA) before offering resistive forces of > 20 pN (**Fig. 1d and 5b**). Assuming a very large loading rate of 200 nm/s will yield a ∼ 0.1 second perturbation of mechanical input (where the force is very low). Based on recent literature, this 0.1 second duration is a sufficient amount time for half of the integrin ɑ_5_β_1_-ligand bonds to dissociate.^28^ Of course some integrins do fully extend the RS probes, leading to activation of their mechanotransduction pathways. Hence, cells cultured on the RS probes display greater markers of mechanotransduction than the irreversible shearing probes.

In conclusion, we have shown that integrin-ligand force duration is highly variable between different processes and cell types. Although the current probe design does not provide a absolute measure of force lifetimes, future iterations of these probes may address this area. Moreover, we show that cell adhesion receptors can detect the molecular force extension curve of their ligand. Previous work focused on cell adhesion receptor sensitivity to only the force threshold of their ligands. Our work indicates that it is not only the force threshold of ligands that controls cell mechanics, but also the unique force-extension curves that these ligands follow. Therefore, further work is necessary to elucidate the exact nature of the receptor-ligand interaction.

## Acknowledgements

This work is supported by NIH NIGMS R01GM124472, NIH NIGMS 1R01GM131099, and NHLBI R01HL142866. R.L.B acknowledges NIH grant no. 3R01GM131099-01S1. H.O. acknowledges the Naito Foundation and the Uehara Memorial Foundation for the research support. A.V.K. acknowledges NIH grant no. F31 F31CA243502. Schemes created with BioRender.com. Mouse platelets were provided by Dr. Wenchun Chen and Prof. Dr. Renhao Li. The authors would like to thank the Emory Mass Spectrometry Center and Dr. Fred Strobel for ESI-MS measurement and Emory University NMR Research Center for 1H NMR measurement.

